# Low convergent validity of [_11_C]raclopride binding in extrastriatal brain regions: a PET study of within-subject correlations with [_11_C]FLB 457

**DOI:** 10.1101/2020.06.01.127027

**Authors:** Tove Freiburghaus, Jonas E. Svensson, Granville J. Matheson, Pontus Plavén-Sigray, Johan Lundberg, Lars Farde, Simon Cervenka

**Author notes:** Corresponding author: Tove Freiburghaus, Centre for Psychiatry Research, Department of Clinical Neuroscience, Karolinska Institutet and Stockholm Health Care Services, Region Stockholm, Stockholm, Sweden, SE-171 76 Stockholm, Sweden.

## Abstract

Dopamine D2 receptors (D2-R) in extrastriatal brain regions are of high interest for research in a wide range of psychiatric and neurologic disorders. Pharmacological competition studies and test-retest experiments have shown high validity and reliability of the positron emission tomography (PET) radioligand [_11_C]FLB 457 for D2-R quantification in extrastriatal brain regions. However, this radioligand is not available at most research centres. Instead, the medium affinity radioligand [_11_C]raclopride, which has been extensively validated for quantification of D2-R in the high-density region striatum, has been applied also in studies on extrastriatal D2-R. Recently, the validity of this approach has been questioned by observations of low occupancy of [_11_C]raclopride in extrastriatal regions in a pharmacological competition study. Here, we utilise a data set of 16 healthy control subjects examined with both [_11_C]raclopride and [_11_C]FLB 457 to assess the correlation in binding potential (BP_ND_) in extrastriatal brain regions. BPND was quantified using the simplified reference tissue model with cerebellum as reference region. The rank order of mean regional BPND values were similar for both radioligands, and corresponded to previously reported data, both post-mortem and using PET. Nevertheless, weak to moderate within-subject correlations were observed between [_11_C]raclopride and [_11_C]FLB 457 BP_ND_ extrastriatally (Pearson’s R: 0.30 - 0.56), in contrast to very strong correlations between repeated [_11_C]FLB 457 measurements (Pearson’s R: 0.82 - 0.98). These results are likely related to low signal to noise ratio of [_11_C]raclopride in extrastriatal brain regions, and further strengthen the recommendation that extrastriatal D2-R measures obtained with [_11_C]raclopride should be interpreted with caution.

## Introduction

Of the dopamine receptor subtypes, the dopamine D2-receptor (D2-R) has been of central interest in research on many neurological and psychiatric disorders. For instance, early positron emission tomography (PET) studies on striatal brain regions using the medium affinity D2-R radioligand [_11_C]raclopride (K_d_ = 1.2 nM) has provided crucial knowledge on the pharmacological properties of antipsychotic drugs (Farde, Hall, Ehrin, & Sedvall, 1986; Farde et al., 1992; Kapur, Remington, Zipursky, Wilson, & Houle, 1995; Nord & Farde, 2011; Nordström et al., 1993), in addition to demonstrating slightly higher striatal D2-R in patients with schizophrenia compared to healthy controls (Howes et al., 2012). More recently, an involvement in the pathophysiology of neurologic and psychiatric disorders has been suggested also for D2-R in extrastriatal brain regions. However, quantification of extrastriatal D2-R is challenging due to the much lower D2-R density, ranging from 1-30% to that of striatum (Hall et al., 1996).

To quantify D2-R binding in low-density extrastriatal regions, a series of radioligands with high affinity have been developed for both autoradiography and molecular imaging use (de Paulis, 2003). One of those is [_11_C]FLB 457 with the very high affinity of K_d_ = 0.02 nM (Halldin et al., 1995). Occupancy and test-retest PET experiments have shown high validity and reliability, respectively, and the radioligand is therefore well suited for extrastriatal D2-R measurements (Farde et al., 1997; Halldin et al., 1995; Narendran, Himes, & Mason, 2013; Narendran, Mason, Chen, et al., 2011; Narendran, Mason, May, et al., 2011; Sudo et al., 2001; Suhara et al., 1999; Vilkman et al., 2000). The synthesis of [_11_C]FLB 457 is, however, technically demanding since high specific radioactivity is required (Halldin et al., 1995; Olsson, Halldin, & Farde, 2004). Additionally, a limitation of high affinity radioligands such as [_11_C]FLB 457 and [_18_F]fallypride (K_d_ = 0.2 nM) (Slifstein et al., 2004) is that accurate quantification of D2-R in the high-density region striatum is rendered either impossible or very impractical: [_11_C]FLB does not reach equilibrium within feasible scanning durations for carbon-11, and [_18_F]fallypride requires 3-4 hours of measurement. (Christian, Narayanan, Shi, & Mukherjee, 2000; Mukherjee et al., 2002; Slifstein et al., 2004).

Given the drawbacks of very high-affinity D2-R radioligands, some research centres have explored the possibility of using [_11_C]raclopride for measuring D2-R also outside of striatum. To date, several such studies have been conducted in patients with Huntington’s disease (Pavese et al., 2003), schizophrenia (Talvik et al., 2006) and major depression (Jussi Hirvonen et al., 2008) as well as in response to methylphenidate in cocaine addiction (Volkow et al., 1997). Additionally, several studies on extrastriatal D2-R have been conducted in healthy individuals in relation to tetrahydrocannabinol effects (Stokes et al., 2010), physical activity and memory (Köhncke et al., 2018; Salami et al., 2019). Studies showing high test-retest reliability have been purported to support this extended use of [_11_C]raclopride (Alakurtti et al., 2015; J. Hirvonen et al., 2003; Karalija et al., 2019a), although conflicting data exists (Mawlawi et al., 2001; Svensson et al., 2019).

However, in addition to reliability, another necessary step for evaluating the suitability of a radioligand is to assess the validity of obtained outcome measures, i.e. if it measures what we expect it to be measuring. This is commonly tested by assessing specific binding in pharmacological competition (occupancy) studies. For [_11_C]FLB 457 such studies using aripiprazole and haloperidol showed significant displacement in all cortical ROIs (Narendran, Mason, Chen, et al., 2011) as well as in thalamus and temporal cortex (Farde et al., 1997). High specific binding of [_11_C]FLB 457 extrastriatally was also demonstrated in one study where the amount of specific [_11_C]FLB 457 radioactivity was systematically varied (Suhara et al., 1999). With regard to [_11_C]raclopride, a recent study showed no competition for binding in frontal cortex, and the effect in thalamus and temporal cortex was significantly lower than in striatum (Svensson et al., 2019). These findings correspond with a previous occupancy study examining the thalamus, showing low [_11_C]raclopride displacement (Mawlawi et al., 2001). Together, these results suggest that the amount of specific binding of [_11_C]raclopride is very low in some extrastriatal regions, and not quantifiable at all in others.

Another approach to assess the validity of a radioligand is to compare outcome measures with that of already established radioligands (convergent validity). Recently, studies comparing binding values in extrastriatal ROIs from separate cohorts examined with [_11_C]raclopride and [_18_F]Fallypride respectively, showed high correspondence between regional average binding levels (Karalija et al., 2019b; Papenberg et al., 2019). This is in contrast to data from a study showing weak correlations between extrastriatal average [_11_C]raclopride and [_11_C]FLB 457 binding (Egerton et al., 2009). Importantly, between-individual comparisons do not account for individual variability in binding and are therefore not suited for assessing measurement precision. We are not aware of any studies to date reporting between-radioligand correlations for individual ROIs, within the same subjects.

Here, we aimed to evaluate the convergent validity of extrastriatal [_11_C]raclopride, by assessing within-individual correlations between [_11_C]raclopride and [_11_C]FLB 457 binding in sixteen healthy control subjects examined with both radioligands. For reference, results were compared to test-retest correlations for each radioligand, as well as published post-mortem autoradiography data on regional D2-R distribution.

## Materials and methods

### Subjects

PET data from sixteen subjects (8 males, 8 females, 56 ± 8 years old) who participated as healthy controls in a previously published PET study (Cervenka et al., 2006) were re-analysed. The subjects had no history of physical or mental illness as assessed by clinical interview, blood and urine tests, brain MRI and ECG. None of the subjects used nicotine and all were naïve to dopaminergic drugs. The subjects abstained from caffeine during the days of the PET examinations. All subjects gave written informed consent before participation according to the Helsinki declaration. The study was approved by the Ethics and Radiation Safety committees of the Karolinska Institute.

### Study design

All participants underwent PET examinations with both [_11_C]raclopride and [_11_C]FLB 457 in random order. All sixteen subjects were examined in the evening (6-8 p.m.) on two separate days with each radioligand. Additionally, eight participants performed an additional PET examination with [_11_C]raclopride in the morning (10-12 p.m.) on the same day as the evening examination, whereas the other eight performed two PET examinations with [_11_C]FLB 457 in the same manner. All participants thus underwent three PET examinations each. The PET examinations for each individual were performed at a median of 7 days apart (range: 1-27 days).

### MRI and PET examinations

T1-weighted magnetic resonance tomography images (MRI) were obtained using a 1.5T GE Signa system (Milwaukee, WI). Images were reconstructed into a 256 x 256 x 156 matrix with a resolution of 1.02 x 1.02 x 1 mm2. PET examinations were conducted using an ECAT Exact HR system (CTI/ Siemens, Knoxville, TN). To minimize head movement a plaster helmet was customized for each subject and used during both PET and MRI examinations. [_11_C]raclopride and [_11_C]FLB 457 were prepared as described elsewhere (Langer et al., 1999; Sandell et al., 2000).The radioligands were administered as bolus into the antecubital vein. Injected radioactivity and ligand mass was 196 ± 4 MBq and 0.62 ± 0.40 ug for [_11_C]raclopride and 201 ± 37 MBq and 0.58 ± 0.43 ug for [_11_C]FLB 457. Radioactivity was measured in the brain during 51 minutes for [_11_C]raclopride and 87 minutes for [_11_C]FLB 457. The reconstructed volume was displayed as 47 horizontal sections with a center-to-center distance of 3.125 mm and a pixel size of 2.02 x 2.02 mm2.

### Preprocessing and ROI definition

PET images were corrected for head motion using a between-frame-correction realignment procedure (Schain et al., 2012). MR images were reoriented to the AC-PC (anterior and posterior commissure) plane. Freesurfer (version 6.0, http://surfer.nmr.mgh.harvard.edu/) was used to delineate regions of interest (ROIs) on the MRIs. Eight ROIs were chosen: occipital cortex, frontal cortex, temporal cortex, hippocampus, thalamus, amygdala, caudate and putamen, based on relevance for psychiatric and neurologic disorders as well as for the purpose of comparison with previous studies reporting on extrastriatal D2-receptors using [_11_C]raclopride (Alakurtti et al., 2015; J. Hirvonen et al., 2003; Karalija et al., 2019; Svensson et al., 2019). Cerebellar cortical grey matter was used as reference region. To avoid partial volume effects and contamination from neighbouring regions in the reference region, the cerebellum was trimmed in an automated process and included voxels behind and below the posterior tip of the 4th ventricle, above the lowest plane of pons, laterally of the left- and rightmost point of the 4th ventricle (as described by (Svensson et al., 2019)). Using SPM12 (Wellcome Department of Cognitive Neurology, University College, London, UK), the MR image was coregistered to summed PET-images for each examination. The resulting coregistration parameters were used to apply ROIs to the dynamic PET data, to generate time activity curves (TACs) of mean radioactivity within each ROI for each frame.

### Quantification of radioligand binding

Kinetic analysis was performed on TACs using the simplified reference tissue model (SRTM) with cerebellar cortex as reference region. The outcome measure was non-displaceable binding potential (BP_ND_). BP_ND_ is the ratio at equilibrium of specifically bound radioligand to that of nondisplaceable radioligand in tissue and is hence theoretically proportional to the amount of available D2-R (Innis et al., 2007). The SRTM has been validated for both [_11_C]raclopride and [_11_C]FLB 457 (Lammertsma AA & Hume SP, 1996; Olsson, Halldin, Swahn, & Farde, 1999).

### Statistical analysis

Rank order of [_11_C]raclopride and [_11_C]FLB 457 regional average BP_ND_ were compared to post-mortem data from Hall et al (1996) data. For comparison to previously published reports (Karalija et al., 2019; Papenberg et al., 2019), we calculated Pearson’s correlations between average [_11_C]raclopride and [_11_C]FLB 457 BP_ND_ values for each ROI. Pearson’s correlations were subsequently performed within subjects, using individual BP_ND_ values in all ROIs. For comparative purposes, a within subject, within radioligand correlation of BP_ND_ from repeated measurements was calculated for both radioligands. All analyses were performed using R (version 3.5.3, *Great Truth*).

## Results

Descriptive data for [_11_C]raclopride and [_11_C]FLB 457 BP_ND_ in extrastriatal ROIs are shown in Table 1. The regional average BP_ND_ values of the evening PET examinations for each radioligand showed good rank-order agreement to published post-mortem autoradiography data using epidepride (Hall et al., 1996), indicating occipital cortex as the region with the lowest level of binding and the putamen as the region with the highest. The correlation in regional average [_11_C]raclopride and [_11_C]FLB 457 BP_ND_ showed a similar pattern as for previous studies using [_18_F]fallypride (Karalija et al., 2019; Papenberg et al., 2019) (Pearson’s R = 0.97) (Supplementary figure 1).

**Table 1.** Rank-order of D2-receptor density from post-mortem autoradiography, and in vivo [_11_C]raclopride and [_11_C]FLB 457 binding values. The data in the second column is retrieved from Hall et al., (Hall et al., 1996) and describes the level of binding in each ROI relative to the level of binding in the putamen. ROI = Region of interest, OC = occipital cortex, FC = frontal cortex, TC = temporal cortex, HIP = hippocampus, THA = thalamus, AMG = amygdala, CAU = caudate, PUT = putamen, BPND = non-displaceable binding potential, sd = standard deviation.

| ROI | Post-mortem data: binding<br>relative to putamen (%) | $\text{BP}_{\text{ND}}$ [$^{11}\text{C}$]raclopride<br>mean (sd) | $\text{BP}_{\text{ND}}$ [$^{11}\text{C}$]FLB-457<br>mean (sd) |
| --- | --- | --- | --- |
| OC | 0.07 | 0.17 (0.06) | 0.43 (0.22) |
| FC | 0.17 | 0.12 (0.05) | 0.48 (0.18) |
| TC | 1.48 | 0.22 (0.06) | 1.10 (0.32) |
| HIP | 2.58 | 0.24 (0.08) | 0.95 (0.24) |
| THA | 3.40 | 0.35 (0.08) | 2.57 (0.54) |
| AMG | 13.80 | 0.36 (0.09) | 2.50 (0.58) |
| CAU | 87.30 | 2.01 (0.23) | NA |
| PUT | 100.00 | 2.79 (0.26) | NA |

Subsequently, we directly compared [_11_C]raclopride and [_11_C]FLB 457 BP_ND_ in individual extrastriatal ROIs, within subjects. Weak to moderate correlations were observed in all extrastriatal ROIs (Figure 1). The R-values ranged from 0.30 to 0.56 (Table 2). The highest correlations were obtained in the amygdala (Pearson’s R: 0.56), the thalamus (Pearson’s R: 0.50) and temporal cortex (Pearson’s R: 0.46). Hence, the explained variance R2 for these regions ranged from 9 to 31%.

**Figure 1.**
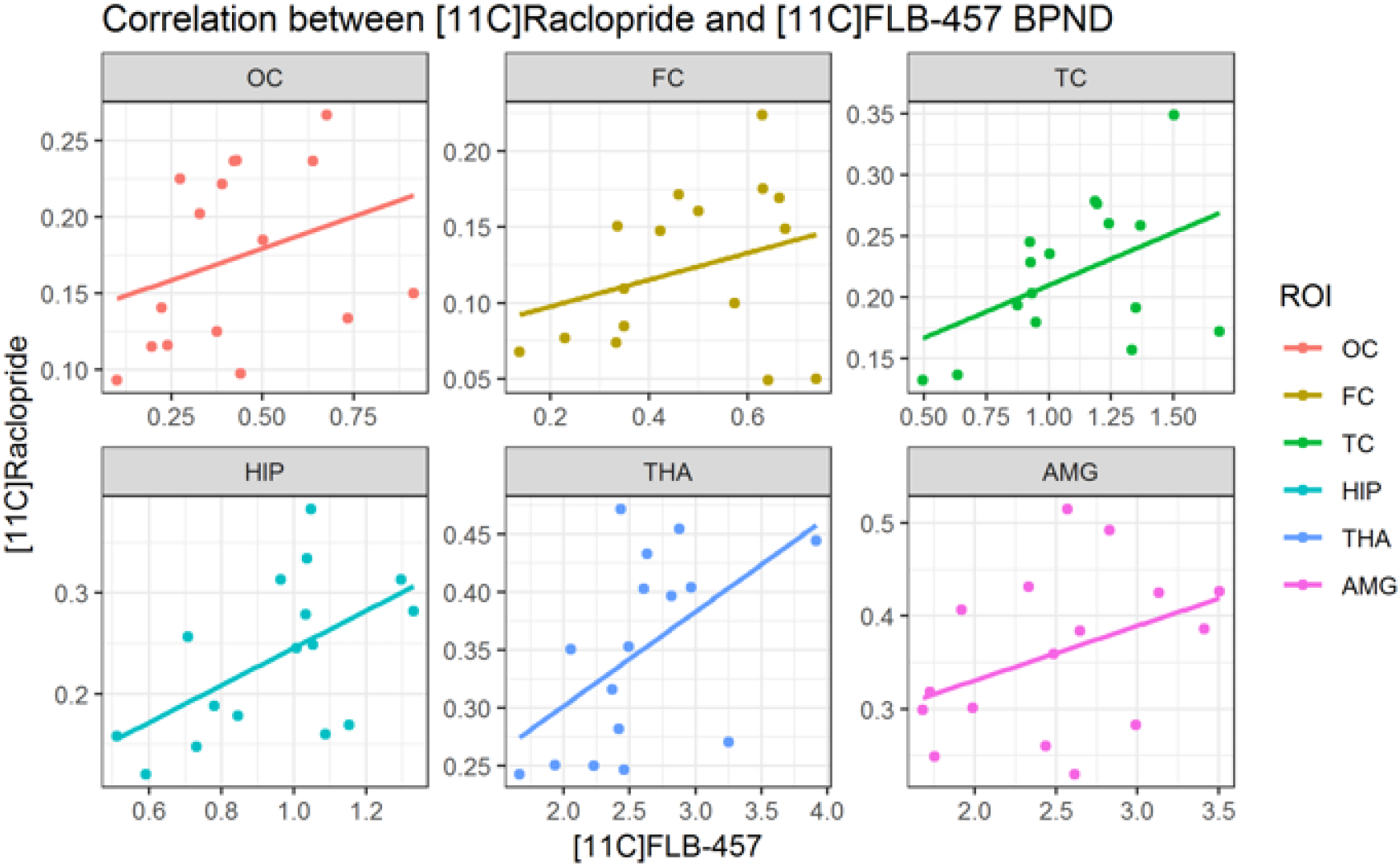
Scatter plots of the relationship between [_11_C]raclopride and [_11_C]FLB 457 BP_ND_ in individual extrastriatal ROIs. OC = occipital cortex, FC = frontal cortex, TC = temporal cortex, HIP = hippocampus, THA = thalamus, AMG = amygdala.

**Table 2.** Table with correlation coefficients and confidence intervals: [_11_C]raclopride to [_11_C]FLB 457 (evening PET-examinations); [_11_C]FLB 457 to [_11_C]FLB 457(repeated measurements); [_11_C]raclopride to [_11_C]raclopride (repeated measurements). 95% CI = confidence interval. ROI = region of interest. OC = occipital cortex, FC = frontal cortex, TC = temporal cortex, HIP = hippocampus, THA = thalamus, AMG = amygdala.

| ROI | [11C]raclopride –<br>[11C]FLB 457<br>Pearson's R (95% CI) | [11C]FLB 457 repeated<br>measurements<br>Pearson's R (95% CI) | [11C]raclopride repeated<br>measurements<br>Pearson's R (95% CI) |
| --- | --- | --- | --- |
| OC | 0.30 (-0.23 – 0.69) | 0.94 (0.71 – 0.99) | 0.04 (-0.68 – 0.72) |
| FC | 0.31 (-0.22 – 0.70) | 0.97 (0.82 – 0.99) | 0.51 (-0.30 – 0.89) |
| TC | 0.46 (-0.05 – 0.78) | 0.96 (0.78 – 0.99) | 0.28 (-0.53 – 0.82) |
| HIP | 0.39 ( -0.13 – 0.74) | 0.82 (0.27 – 0.97) | 0.35 (-0.48 – 0.85) |
| THA | 0.50 (0.00 – 0.80) | 0.98 (0.89 – 1.00) | 0.46 (-0.37 – 0.88) |
| AMG | 0.56 (0.09 – 0.83) | 0.89 (0.51 – 0.98) | 0.75 (0.10 – 0.95) |

Within-radioligand correlations for both [_11_C]raclopride and [_11_C]FLB-457 are presented in Figure 2 as well as Table 2. For [_11_C]FLB 457 the average Pearson’s R was 0.93, whereas for [_11_C]raclopride the corresponding average was 0.40. For additional test-retest metrics for [_11_C]raclopride and [_11_C]FLB 457 see supplementary material (Supplementary Table 1 and 2).

**Figure 2.**
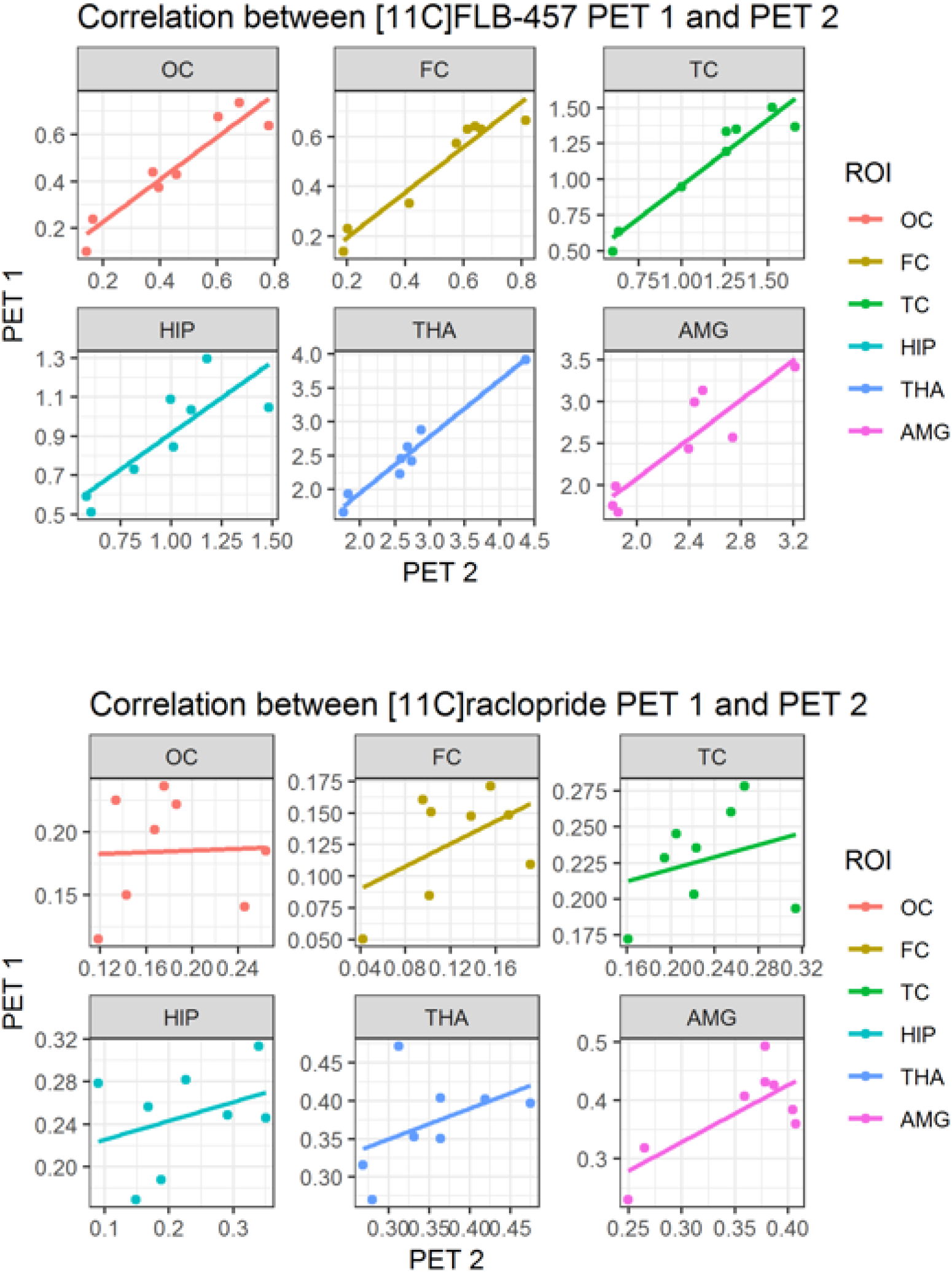
Scatter plot of test-retest reliability for [_11_C]FLB 457 and [_11_C]raclopride BP_ND_ in individual extrastriatal ROIs. OC = occipital cortex, FC = frontal cortex, TC = temporal cortex, HIP = hippocampus, THA = thalamus, AMG = amygdala.

## Discussion

To our knowledge this is the first study to report on within-subject correlations of binding in extrastriatal ROIs between [_11_C]raclopride and a very high affinity D2-R radioligand. Overall, the agreement to [_11_C]FLB 457 BP_ND_ was found to be low, indicating low convergent validity of [_11_C]raclopride for measurements in extrastriatal brain regions. Given that multiple competition and test-retest studies have demonstrated that [_11_C]FLB 457 is a well suited radioligand to index extrastriatal D2-R binding (Farde et al., 1997; Halldin et al., 1995; Narendran et al., 2013; Narendran, Mason, Chen, et al., 2011; Narendran, Mason, May, et al., 2011; Sudo et al., 2001; Suhara et al., 1999; Vilkman et al., 2000), the results imply that the low correlation to [_11_C]raclopride binding is due to low precision of [_11_C]raclopride for extrastriatal D2-R quantification.

Reports of associations in regional average BP_ND_ between [_11_C]raclopride and post-mortem data, as well as between [_11_C]raclopride and the very high affinity radioligand [_18_F]fallypride, have been purported to support the use of [_11_C]raclopride extrastriatally (Karalija et al., 2019; Papenberg et al., 2019), although conflicting data have also been presented (Egerton et al., 2009). In the present study we could confirm a rank order association between regional average [_11_C]raclopride and [_11_C]FLB 457 BP_ND_ as well as with post-mortem data. These results may be interpreted as support for some degree of accuracy (i.e. closeness to the “true” underlying values) of [_11_C]raclopride extrastriatally, but only on a group level. It should be noted that all these analyses include ROIs with markedly different levels of D2-R density, thus forming subgroups which can lead to spurious correlations (Makin & De Xivry, 2019). Importantly, our results clearly show that associations between regional averages do not predict the strength of within-subject correlations. In the present study, the correlation coefficient for extrastriatal regions ranged between 0.30 - 0.56, corresponding to a median R2 of 0.18, which means that only about 18% of the variation in [_11_C]raclopride BP_ND_ is explained by variation in [_11_C]FLB 457 BP_ND_. For two methods purported to measure the same thing, this agreement is very poor. For comparison, the correlation coefficient for repeated [_11_C]FLB 457 measurements was 0.82 - 0.98, corresponding to a median explained variance of 90%. In summary, even if some specific binding may be detectable in extrastriatal regions, such that some degree of accuracy can be claimed based on group data, this does not necessarily indicate adequate precision, and thereby validity, of extrastriatal [_11_C]raclopride measurements of D2-R at the individual level.

A likely factor explaining the observed low precision of [_11_C]raclopride in extrastriatal regions is the very low level of specific binding, leading to a low signal-to-noise ratio. Importantly, an in vitro study with the radioligand [_3_H]raclopride, with the same affinity as [_11_C]raclopride (Hall, Köhler, Gawell, Farde, & Sedvall, 1988), showed no specific binding in amygdala, hippocampus or cerebellum, whereas specific binding in frontal and temporal cortex was much lower compared to putamen and caudate (Hall, Farde, & Sedvall, 1988). Low specific in relation to non-specific binding in extrastriatal regions has also been confirmed in *in vivo* occupancy studies. In two small occupancy studies using haloperidol, in 4/5 subjects only roughly 50% of [_11_C]raclopride BP_ND_ in thalamus corresponded to specific binding (Hirvonen et al., 2003; Mawlawi et al., 2001). More recently, lower occupancy of [_11_C]raclopride was shown by Svensson et al. in a larger competition study (n=9) using quetiapine. Lower occupancy in thalamus compared to striatum was observed in both high and low dose regimens (20% of thalamic BP_ND_ was displaced with doses displacing 50% of BP_ND_ in striatum) (Svensson et al., 2019). Additionally, only 18 % of raclopride binding was occupied in the temporal cortex, whereas in frontal cortical regions and the anterior cingulate no clear reduction was seen in BP_ND_ after administration of quetiapine, suggesting no specific binding in these regions. The corresponding pattern between this competition study and the present results, with numerically lower correlations in frontal cortical regions compared to thalamus and temporal cortex, suggest decreasing validity of [_11_C]raclopride measurements as a function of lower D2-R density.

In addition to validity, reliability provides additional information about a method that can guide its use (Matheson, 2019). Test-retest studies of [_11_C]FLB 457 using SRTM have shown high reliability in cortical brain regions and thalamus (Sudo et al., 2001; Vilkman et al., 2000), a result later replicated in additional extrastriatal brain regions (Narendran et al., 2013; Narendran, Mason, May, et al., 2011). In the present study we confirmed high test-retest reliability of [_11_C]FLB 457. For [_11_C]raclopride, some test-retest studies in extrastriatal regions have shown high reliability (Alakurtti et al., 2015; J. Hirvonen et al., 2003; Karalija et al., 2019a). In contrast, our observations of low to moderate test-retest reliability are in line with results by Mawlawi et al and Svensson et al (Mawlawi et al., 2001; Svensson et al., 2019). The question regarding the reliability of [_11_C]raclopride extrastriatally remains to be resolved. However it should be noted that this is of secondary importance if the validity of measurements is inadequate. I.e. it is of little use to assess the reliability or consistency of extrastriatal D2-R measurements, when we cannot ascertain that BPND is a suitable index of D2-R availability.

One potential explanation that has been proposed for the putative combination of low validity and high reliability of extrastriatal [_11_C]raclopride measurements is that BP_ND_ in these regions may be inflated by a contribution of non-displaceable distribution volume (V_ND_) due to lower non-specific binding in the reference region compared to target regions. Since V_ND_ is assumed to be stable over time, such a contamination effect would lead to over-estimated reliability measures (Mawlawi et al., 2001; Svensson et al., 2019, 2020).

Our results have implications for the evidential value of previous studies reporting on extrastriatal binding measures using [_11_C]raclopride. Even for the thalamus, where we found numerically higher correlations and aforementioned occupancy studies support some degree of specific [_11_C]raclopride binding (Hirvonen et al., 2003; Mawlawi et al., 2001; Svensson et al., 2019), most PET studies have insufficient power to detect anything less than large effect sizes (Svensson et al., 2019, 2020). With low power comes an elevated risk for type II, and potentially also type I errors (Button et al., 2013; Loken & Gelman, 2017). A previous [_11_C]raclopride study from our lab reported lower D2-R BP_ND_ in the right thalamus in 18 drug-naïve patients with schizophrenia compared to 17 control subjects (Talvik et al., 2006). Even though this result is broadly in line with studies using high affinity D2-R radioligands (Graff-Guerrero et al., 2009; Kessler et al., 2009; Lehrer et al., 2010; Suhara et al., 2002; Talvik, Nordström, Olsson, Halldin, & Farde, 2003; Tuppurainen et al., 2006), there is hence reason to question the effect size and conclusion of the study. To avoid problems of both low sensitivity and spurious findings we recommend that future research on extrastriatal D2-R is conducted using high-affinity radioligands only.

There are limitations to this study. Cross-sectional studies have indicated that increasing age is associated with a reduction in D2-R density, with estimates in extrastriatal regions ranging from around 5 – 13 % per decade, as well as lower interindividual differences in BP_ND_ (Inoue et al., 2001; Kaasinen et al., 2000; Rinne et al., 1993). The mean age of the subjects in this study was 56 years. It is conceivable that higher D2-R levels in a younger sample could increase the signal-to-noise ratio for extrastriatal [_11_C]raclopride binding measures. Moreover, the relatively high age of the participants may have contributed to lower ICC values in the test-retest analysis. Furthermore, [_11_C]FLB 457 have a low but detectable level of specific binding in the cerebellum (Narendran, Mason, Chen, et al., 2011; Olsson et al., 2004), although the impact on BP_ND_ in target regions has been shown to be small (Olsson et al., 2004). Finally, the experiments for between-radioligand comparisons were further apart compared to experiments for within-radioligand comparisons which may have contributed to the low correlations between [_11_C]raclopride and [_11_C]FLB 457, however it should be noted that the within-radioligand correlations for [_11_C]raclopride were similarly low, whereas corresponding correlations for [_11_C]FLB 457 were high.

In conclusion, our study adds to recently published data indicating low validity of [_11_C]raclopride binding measures in extrastriatal brain regions. The results have important implications for the interpretation of previously published data and should inform the design of future PET studies of D2-R outside striatal regions, to save time and resources.

## Supplementary material

**Figure 1:**
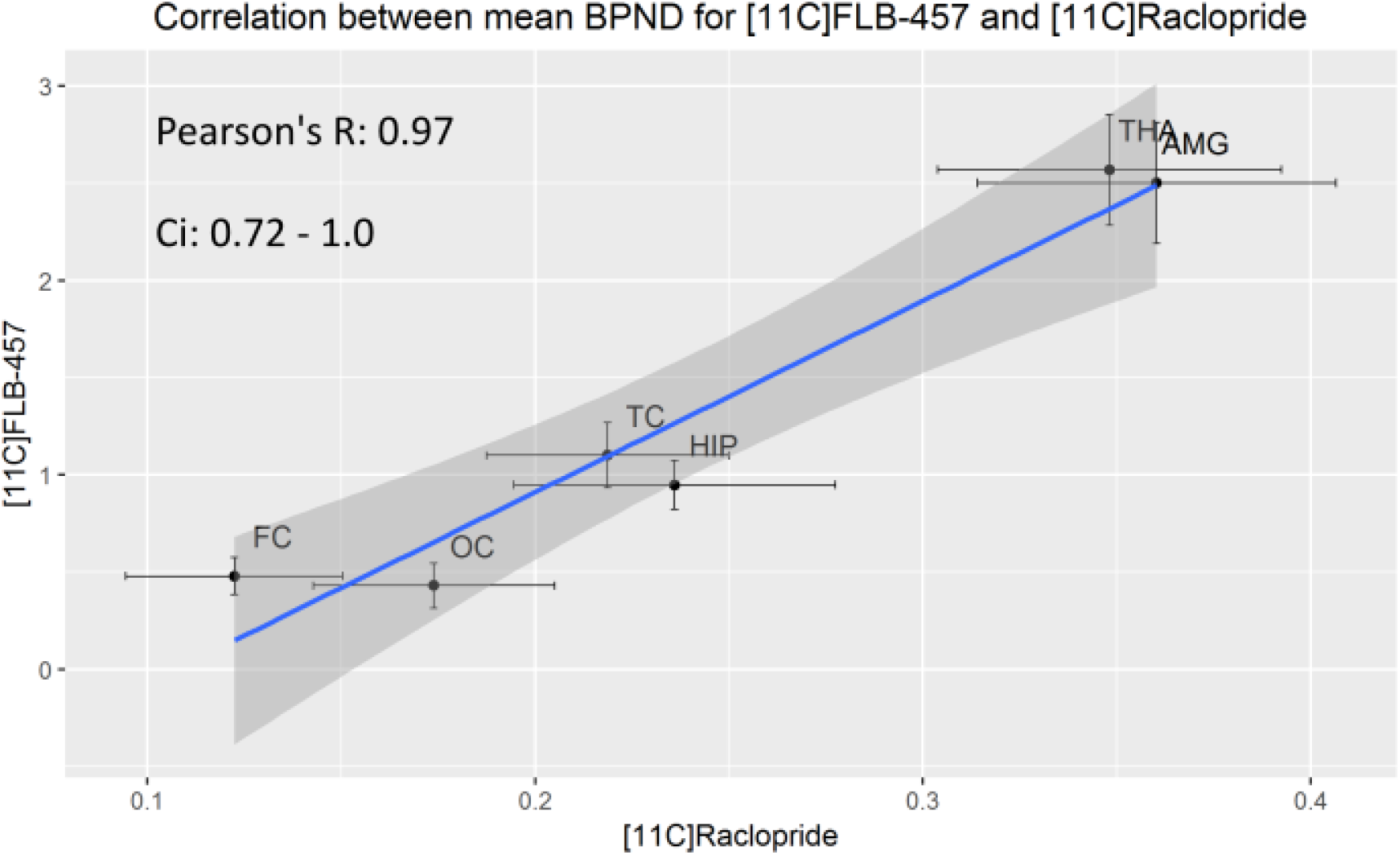
Correlation between regional average [_11_C]raclopride and [_11_C]FLB 457 BP_ND_. Ci: confidence interval. OC = occipital cortex, FC = frontal cortex, TC = temporal cortex, HIP = hippocampus, THA = thalamus, AMG = amygdala. The 95% confidence intervals within each ROI are marked with vertical and horizontal lines.

**Table 1.**
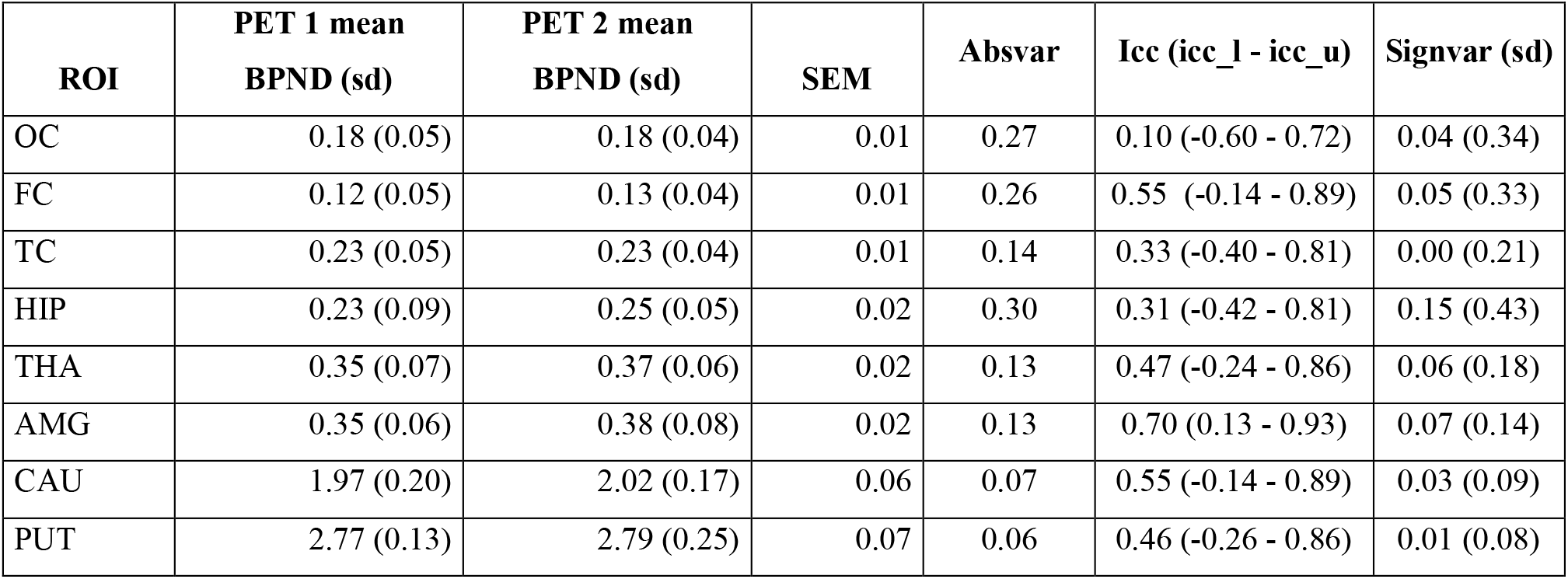
Test-retest for [_11_C]raclopride. Average BPND within each ROI in morning and evening PET experiments (PET 1 and PET 2) as well as standard deviations. SEM = standard error of measurement, absvar = average absolute variability, icc = intraclass correlation coefficient, icc_l and icc_u are the upper and lower bounds of the 95 percent confidence interval, signvar = signed variability (for detecting bias between measurements) normalised to the grand mean.

**Table 2.** Test-retest for [_11_C]FLB Average BPND within each ROI in morning and evening PET experiments (PET 1 and PET 2) as well as standard deviations. SEM = standard error of measurement, absvar = average absolute variability, icc = intraclass correlation coefficient, icc_l and icc_u are the upper and lower bounds of the 95 percent confidence interval, signvar = signed variability (for detecting bias between measurements) normalised to the grand mean.

| **ROI** | **PET 1 mean**<br>**BPND (sd)** | **PET 2 mean**<br>**BPND (sd)** | **SEM** | **Absvar** | **Icc (icc_l - icc_u)** | **Signvar (sd)** |
| --- | --- | --- | --- | --- | --- | --- |
| OC | 0.45 (0.23) | 0.45 (0.22) | 0.05 | 0.17 | 0.95 (0.79 - 0.99) | 0.00 (0.22) |
| FC | 0.51 (0.23) | 0.48 (0.21) | 0.04 | 0.12 | 0.96 (0.80 - 0.99) | -0.08 (0.15) |
| TC | 1.16 (0.39) | 1.10 (0.37) | 0.08 | 0.07 | 0.95 (0.80 - 0.99) | -0.05 (0.09) |
| HIP | 0.97 (0.30) | 0.89 (0.27) | 0.06 | 0.13 | 0.80 (0.34 - 0.96) | -0.08 (0.15) |
| THA | 2.68 (0.80) | 2.52 (0.68) | 0.13 | 0.07 | 0.95 (0.66 - 0.99) | -0.06 (0.07) |
| AMG | 2.35 (0.50) | 2.50 (0.65) | 0.15 | 0.10 | 0.85 (0.46 - 0.97) | 0.05 (0.12) |
| CAU | NA | NA | NA | NA | NA | NA |
| PUT | NA | NA | NA | NA | NA | NA |

## References

Alakurtti, K., Johansson, J. J., Joutsa, J., Laine, M., Bäckman, L., Nyberg, L., & Rinne, J. O. (2015). Long-term test-retest reliability of striatal and extrastriatal dopamine D2/3 receptor binding: Study with [11C]raclopride and high-resolution PET. Journal of Cerebral Blood Flow and Metabolism, 35(7), 1199–1205.

Button, K. S., Ioannidis, J. P. A., Mokrysz, C., Nosek, B. A., Flint, J., Robinson, E. S. J., & Munafò, M. R. (2013). Power failure: Why small sample size undermines the reliability of neuroscience. Nature Reviews Neuroscience, 14(5), 365–376.

Cervenka, S., Pålhagen, S. E., Comley, R. A., Panagiotidis, G., Cselényi, Z., Matthews, J. C., Lai, R. Y., Halldin, C., Farde, L. (2006). Support for dopaminergic hypoactivity in restless legs syndrome: A PET study on D2-receptor binding. Brain, 129(8), 2017–2028.

Christian, B. T., Narayanan, T. K., Shi, B., & Mukherjee, J. (2000). Quantitation of striatal and extrastriatal D-2 dopamine receptors using PET imaging of [18F]fallypride in nonhuman primates. Synapse, 38(1), 71–79.

de Paulis, T. (2003). The discovery of epidepride and its analogs as high-affinity radioligands for imaging extrastriatal dopamine D(2) receptors in human brain. Current Pharmaceutical Design, 9(8), 673–696.

Egerton, A., Mehta, M. A., Montgomery, A. J., Lappin, J. M., Howes, O. D., Reeves, S. J., Cunningham, V. J., Grasby, P. M. (2009). The dopaminergic basis of human behaviors: A review of molecular imaging studies. Neuroscience and Biobehavioral Reviews, 33(7), 1109–1132.

Farde, L., Hall, H., Ehrin, E., & Sedvall, G. (1986). Quantitative analysis of D2 dopamie receptor binding in the living human brain by PET. Science, 231(4735), 258–261.

Farde, L., Nordström, A.-L., Wiesel, F.-A., Pauli, S., Halldin, C., & Sedvall, G. (1992). Positron Emission Tomographic Analysis of Central D1 and D2 Dopamine Receptor Occupancy in Patients Treated With Classical Neuroleptics and Clozapine Relation to. Arch Gen Psychiatry, 49(7), 538–544.

Farde, L., Suhara, T., Nyberg, S., Karlsson, P., Nakashima, Y., Hietala, J., & Halldin, C. (1997). A PET-study of [11C]FLB 457 binding to extrastriatal D2-dopamine receptors in healthy subjects and antipsychotic drug-treated patients. Psychopharmacology, 133(4), 396–404.

Graff-guerrero, A., Mizrahi, R., Agid, O., Marcon, H., Barsoum, P., & Rusjan, P. (2009). The Dopamine D 2 Receptors in High-Affinity State and D 3 Receptors in Schizophrenia: A Clinical [11 C] - (+) -PHNO PET Study. Neuropsychopharmacology, 34(4), 1078–1086.

Hall, H., Farde, L., Halldin, C., Hurd, Y. L., Pauli, S., & Sedvall, G. (1996). Autoradiographic localization of extrastriatal D2-dopamine receptors in the human brain using [125I]epidepride. Synapse, 23(2), 115–123.

Hall, H., Köhler, C., Gawell, L., Farde, L., & Sedvall, G. (1988). Raclopride, a new selective ligand for the dopamine-D2 receptors. Progress in Neuropsychopharmacology and Biological Psychiatry, 12(5), 559–568.

Halldin, C., Farde, L., Hogberg, T., Mohell, N., Hall, H., Suhara, T., Karlsson, P., Nakashima, Y., Swahn, C. G. (1995). Carbon-11-FLB 457: A radioligand for extrastriatal D2 dopamine receptors. Journal of Nuclear Medicine, 36(7), 1275–1281.

Hirvonen, J., Aalto, S., Lumme, V., Någren, K., Kajander, J., Vilkman, H., Hagelberg, N., Oikonen, V., Hietala, J. (2003). Measurement of striatal and thalamic dopamine d2 receptor binding with c-raclopride. Nuclear Medicine Communications, 24(12), 1207–1214.

Hirvonen, J., Karlsson, H., Kajander, J., Markkula, J., Rasi-Hakala, H., Någren, K., Salminen, J. K., Hietala, J. (2008). Striatal dopamine D2 receptors in medication-naive patients with major depressive disorder as assessed with [11C]raclopride PET. Psychopharmacology, 197(4), 581–590.

Howes, O. D., Kambeitz, J., Kim, E., Stahl, D., Slifstein, M., Abi-Dargham, A., & Kapur, S. (2012). The nature of dopamine dysfunction in schizophrenia and what this means for treatment: Meta-analysisof imaging studies. Archives of General Psychiatry, 69(8), 776–786.

Innis, R. B., Cunningham, V. J., Delforge, J., Fujita, M., Gjedde, A., Gunn, R. N., Holden, J., Houle, S., Hoang, S. C., Ichise, M., Iida, H., Ito, H., Kimura, Y., Koeppe, R. A., Knudsen, G. M., Knuuti, J., Lamertsmaa, A. A., Laruelle, M., Logan, J., Maguire, R. P., Mintun, M. A., Morris, E. D., Parsey, R., Price, J. C., Slifstein, M., Sossi, V., Suhara, T., Votaw, J. R., Wong, D. F., Carson, R. E. (2007). Consensus nomenclature for in vivo imaging of reversibly binding radioligands. Journal of Cerebral Blood Flow and Metabolism, 27(9), 1533–1539.

Inoue, M., Suhara, T., Sudo, Y., Okubo, Y., Yasuno, F., Kishimoto, T., Yoshikawa, K., Tanada, S. (2001). Age-related reduction of extrastriatal dopamine D2 receptor measured by PET. Life Sciences, 69(9), 1079–1084.

Kaasinen, V., Vilkman, H., Hietala, J., Någren, K., Helenius, H., Olsson, H., Farde, L., Rinne, J. O. (2000). Age-related dopamine D2/D3 receptor loss in extrastriatal regions of the human brain. Neurobiology of Aging, 21(5), 683–688.

Kapur, S., Remington, G., Zipursky, R. B., Wilson, A. A., & Houle, S. (1995). The D2 dopamine receptor occupancy of risperidone and its relationship to extrapyramidal symptoms: A pet study. Life Sciences, 57(10).

Karalija, N., Jonassson, L., Johansson, J., Papenberg, G., Salami, A., Andersson, M., Katrine, R., Nyberg, L., Boraxbekk, C. J. (2019). High long-term test-retest reliability for extrastriatal 11C-raclopride binding in healthy older adults. Journal of Cerebral Blood Flow and Metabolism.

Kessler, R. M., Woodward, N. D., Riccardi, P., Li, R., Ansari, M. S., Anderson, S., Dawant, B., Zald, D., Meltzer, H. Y. (2009). Dopamine D2 Receptor Levels in Striatum, Thalamus, Substantia Nigra, Limbic Regions, and Cortex in Schizophrenic Subjects. Biological Psychiatry, 65(12), 1024–1031.

Köhncke, Y., Papenberg, G., Jonasson, L., Karalija, N., Wåhlin, A., Salami, A., Andersson, M., Axelsson, J. E., Nyberg, L., Riklund, K., Bäckman, L., Lindenberger, U., Lövdén, M. (2018). Self-rated intensity of habitual physical activities is positively associated with dopamine D2/3 receptor availability and cognition. NeuroImage, 181(July), 605–616.

Lammertsma AA, & Hume SP. (1996). Simplified reference tissue model for PET receptor studies. Neuroimage, 4(4), 153–158.

Langer, O., Någren, K., Dolle, F., Lundkvist, C., Sandell, J., Swahn, C.-G., Vaufrey, F., Crouzel, C., Maziere, B., Halldin, C. (1999). Precursor synthesis and radiolabelling of the dopamine D2 receptor ligand [11C]raclopride from [11C]methyl triflate. Journal of Labelled Compounds and Radiopharmaceuticals, 42(12), 1183–1193.

Lehrer, D. S., Christian, B. T., Kirbas, C., Chiang, M., Sidhu, S., Short, H., Wang, B., Shi, B., Chu, K., Merrill, B., Buchsbaum, M. S. (2010). F-Fallypride binding potential in patients with schizophrenia compared to healthy controls. Schizophrenia Research, 122(1–3), 43–52.

Loken, E., & Gelman, A. (2017). Measurement error and the replication crisis. Science, 355(6325), 584–585.

Makin, T. R., & De Xivry, J. J. O. (2019). Ten common statistical mistakes to watch out for when writing or reviewing a manuscript. ELife, 8, 1–13.

Matheson, G. J. (2019). We need to talk about reliability: making better use of test-retest studies for study design and interpretation. PeerJ, 7, e6918.

Mawlawi, O., Martinez, D., Slifstein, M., Broft, A., Chatterjee, R., Hwang, D. R., Huang, Y., Simpson, N., Ngo, K., Van Heertum, R., Laruelle, M. (2001). Imaging human mesolimbic dopamine transmission with positron emission tomography: I. Accuracy and precision of D2 receptor parameter measurements in ventral striatum. Journal of Cerebral Blood Flow and Metabolism, 21(9), 1034–1057.

Mukherjee, J., Christian, B. T., Dunigan, K. A., Shi, B., Narayanan, T. K., Satter, M., & Mantil, J. (2002). Brain imaging of 18F-fallypride in normal volunteers: Blood analysis, distribution, test-retest studies, and preliminary assessment of sensitivity to aging effects on dopamine D-2/D-3 receptors. Synapse, 46(3), 170–188.

Narendran, R., Himes, M., & Mason, N. S. (2013). Reproducibility of post-amphetamine [11C]FLB 457 binding to cortical D2/3 receptors. PloS One, 8(9), e76905.

Narendran, R., Mason, N. S., Chen, C., Himes, M., Keating, P., May, M. A., Rabiner, E. A., Laruelle, M., Mathis, C. A., Frankle, G. (2011). Evaluation of dopamine D2/3 specific binding in the cerebellum for the PET radiotracer [11C]FLB 457: implications for measuring cortical dopamine release. Synapse, 65(10), 991–997.

Narendran, R., Mason, N. S., May, M. A., Chen, C.-M., Kendro, S., Ridler, K., Rabiner, E. A., Laruelle, M., Mathis, C. A., Frankle, W. G. (2011). Positron emission tomography imaging of dopamine D2/3 receptors in the human cortex with [11C]FLB 457: reproducibility studies. Synapse (New York, N.Y.), 65(1), 35–40.

Nord, M., & Farde, L. (2011). Antipsychotic Occupancy of Dopamine Receptors in Schizophrenia. CNS Neuroscience and Therapeutics, 17(2), 97–103.

Nordström, A. L., Farde, L., Wiesel, F. A., Forslund, K., Pauli, S., Halldin, C., & Uppfeldt, G. (1993). Central D2-dopamine receptor occupancy in relation to antipsychotic drug effects: A double-blind PET study of schizophrenic patients. Biological Psychiatry, 33(4), 227–235.

Olsson, H., Halldin, C., & Farde, L. (2004). Differentiation of extrastriatal dopamine D2 receptor density and affinity in the human brain using PET. NeuroImage, 22(2), 794–803.

Olsson, H., Halldin, C., Swahn, C., & Farde, L. (1999). ll Quantification of [C] FLB 457 Binding to Extrastriatal Dopamine Receptors in the Human Brain. Blood, 19(10), 1164–1173.

Papenberg, G., Jonasson, L., Karalija, N., Johansson, J., Köhncke, Y., Salami, A., Andersson, M., Axelsson, J., Wåhlin, A., Riklund, K., Lindenberger, U., Lövdén, M., Nyberg, L., Bäckman, L. (2019). Mapping the landscape of human dopamine D2/3 receptors with [11C]raclopride. Brain Structure and Function, 224(8), 2871–2882.

Pavese, N., Andrews, T. C., Brooks, D. J., Ho, A. K., Rosser, A. E., Barker, R. A., Robins, T. W., Sahakian, B. J., Dunnett., S. B., Piccini, P. (2003). Progressive striatal and cortical dopamine receptor dysfunction in Huntington’s disease: A pet study. Brain, 126(5), 1127–1135.

Rinne, J. O., Hietala, J., Ruotsalainen, U., Säkö, E., Laihinen, A., Någren, K., Lehikoinen, P., Oikonen, V., Syvälahti, E. (1993). Decrease in human striatal dopamine D2 receptor density with age: A PET study with [11C]raclopride. In Journal of Cerebral Blood Flow and Metabolism (Vol. 13).

Salami, A., Garrett, D. D., Wåhlin, A., Rieckmann, A., Papenberg, G., Karalija, N., Jonasson, L., Andersson, M., Axelsson, J., Johansson, J., Riklund, K., Lövdén, M., Lindenberger, U., Bäckman, L., Nyberg, L. (2019). Dopamine D 2/3 binding potential modulates neural signatures of working memory in a load-dependent fashion. Journal of Neuroscience, 39(3), 537–547.

Sandell, J., Langer, O., Larsen, P., Dolle, F., Vaufrey, F., Demphel, S., Crouzel, C., Halldin, C. (2000). Improved specific radioactivity of the PET radioligand [11C]FLB 457 by use of the GE medical systems PETtrace MeI microlab. Journal of Labelled Compounds and Radiopharmaceuticals, 43(4), 331–338.

Schain, M., Tóth, M., Cselényi, Z., Stenkrona, P., Halldin, C., Farde, L., & Varrone, A. (2012). Quantification of serotonin transporter availability with [11C]MADAM - A comparison between the ECAT HRRT and HR systems. NeuroImage, 60(1), 800–807.

Slifstein, M., Hwang, D. R., Huang, Y., Guo, N. N., Sudo, Y., Narendran, R., Talbot, P., Laruele, M. (2004). In vivo affinity of [18F]fallypride for striatal and extrastriatal dopamine D2 receptors in nonhuman primates. Psychopharmacology, 175(3), 274–286.

Stokes, P. R. A., Egerton, A., Watson, B., Reid, A., Breen, G., Lingford-Hughes, A., Nutt, D. J., Mehta, M. A. (2010). Significant decreases in frontal and temporal [11C]-raclopride binding after THC challenge. NeuroImage, 52(4), 1521–1527.

Sudo, Y., Sudo, Y., Suhara, T., Suhara, T., Inoue, M., Ito, H., Suzuki, K., Saijo, T., Halldin, C., Farde, L. (2001). Reproducibility of [11 c]flb 457 binding in extrastriatal regions. Nuclear Medicine Communications, 22(11), 1215–1221.

Suhara, T., Okubo, Y., Yasuno, F., Sudo, Y., Inoue, M., Ichimiya, T., Nakashima, Y., Nakayama, K., Tanada, S., Suzuki, K., Halldin, C., Farde, L. (2002). Decreased dopamine D2 receptor binding in the anterior cingulate cortex in schizophrenia. Archives of General Psychiatry, 59(1), 25–30.

Suhara, T., Sudo, Y., Okauchi, T., Maeda, J., Kawabe, K., Suzuki, K., Okubo, Y., Nakashima, Y., Ito, H., Tanada, S., Halldin, C., Farde, L. (1999). Extrastriatal dopamine D2 receptor density and affinity in the human brain measured by 3D PET. International Journal of Neuropsychopharmacology, 2(2), 73–82.

Svensson, J. E., Schain, M., Plavén-Sigray, P., Cervenka, S., Tiger, M., Nord, M., Halldin, C., Farde, L., Lundberg, J. (2019). Validity and reliability of extrastriatal [11C]raclopride binding quantification in the living human brain. NeuroImage, 202, 116143.

Svensson, J. E., Schain, M., Plavén-Sigray, P., Cervenka, S., Tiger, M., Nord, M., Halldin, C., Farde, L., Lundberg, J. (2020). In response to the letter “ [11 C]raclopride and extrastriatal binding to D2/3 receptors” by Dr. Heiko Backes. NeuroImage, 207, 116371.

Takahashi, K., Mizuno, K., Sasaki, A. T., Wada, Y., Tanaka, M., Ishii, A., Tajima, K., Tsuyuguchi, N., Watanabe, K., Zeki, S., Watanabe, Y. (2015). Imaging the passionate stage of romantic love by dopamine dynamics. Frontiers in Human Neuroscience, 9(191).

Talvik, M., Nordström, A. L., Okubo, Y., Olsson, H., Borg, J., Halldin, C., & Farde, L. (2006). Dopamine D2 receptor binding in drug-naïve patients with schizophrenia examined with raclopride-C11 and positron emission tomography. Psychiatry Research - Neuroimaging, 148(2–3), 165–173.

Talvik, M., Nordström, A. L., Olsson, H., Halldin, C., & Farde, L. (2003). Decreased thalamic D2/D3 receptor binding in drug-naive patients with schizophrenia: A PET study with [11C]FLB 457. International Journal of Neuropsychopharmacology, 6(4), 361–370.

Tuppurainen, H., Kuikka, J. T., Laakso, M. P., Viinamäki, H., Husso, M., & Tiihonen, J. (2006). Midbrain dopamine D 2/3 receptor binding in schizophrenia. Eur Arch Psychiatry Clin Neurosci, 382–387.

Vilkman, H., Kajander, J., Någren, K., Oikonen, V., Syvälahti, E., & Hietala, J. (2000). Measurement of extrastriatal D2-like receptor binding with [11C]FLB 457 - A test-retest analysis. European Journal of Nuclear Medicine, 27(11), 1666–1673.

Volkow, N. D., Wang, G. J., Fowler, J. S., Logan, J., Gatley, S. J., Hitzemann, R., Chen, A. D., Dewey, S. L., Pappas, N. (1997). Decreased striatal dopaminergic responsiveness in detoxified cocaine-dependent subjects. Nature, 386(6627), 830–833.

